# Cooperation Enhances Robustness of Coexistence in Spatially Structured Consortia

**DOI:** 10.1101/472811

**Authors:** Xinying Ren, Richard M. Murray

## Abstract

Designing synthetic microbial consortia is an emerging area in synthetic biology and a major goal is to realize stable and robust coexistence of multiple species. Co-operation and competition are fundamental intra/interspecies interactions that shape population level behaviors, yet it is not well-understood how these interactions affect the stability and robustness of coexistence. In this paper, we show that communities with cooperative interactions are more robust to population disturbance, e.g., depletion by antibiotics, by forming intermixed spatial patterns. Meanwhile, competition leads to population spatial heterogeneity and more fragile coexistence in communities. Using reaction-diffusion and nonlocal PDE models and simulations of a two-species *E. coli* consortium, we demonstrate that cooperation is more beneficial than competition in maintaining coexistence in spatially structured consortia, but not in well-mixed environments. This also suggests a trade-off between constructing heterogeneous communities with localized functions and maintaining robust coexistence. The results provide general strategies for engineering spatially structured consortia by designing interspecies interactions and suggest the importance of cooperation for biodiversity in microbial community.

## I. INTRODUCTION

Microbial consortia exist in all natural environments, such as mammalian guts [1], foods [2], soils [3], water bodies and wastes [4]. Multiple species coexist in consortia and the interactions among species play a key role in their survival [5]. Compared with monocultures, consortia contain more diverse structures and functions to promote stability and robustness to fluctuations in environments. Inspired by the enhanced performance in productivity, efficiency and robustness of natural microbial consortia, researchers have started to design synthetic consortia and regulate population level behaviors of multi-species to achieve complex tasks. By carefully engineering intra/interspecies communication pathways, dividing metabolic labor and assembling functional modules across mixed populations [6], [7], the synthetic consortia can perform multiple tasks or functions with multiple steps and overcome limitations of genetic circuits in single cells.

Recent synthetic consortia designs include the predatorprey system [8], the rock-paper-scissors system [9], an emergent oscillator in two species [10], a toggle switch in two species [11], feedback controllers on population size [12], three- and four-strain ecosystems by social interaction programming [13], etc. Researchers focus on engineering intercellular communication via quorum sensing (QS) signals [14], essential metabolites exchange [15], or secreted enzymes [16], and compartmentalizing functional modules such as biosynthesis of desired chemical products across microbial populations. To ensure the functionality of the consortia, it is important to maintain a stable coexistence of multiple species and robust cell-cell interactions.

The stability and adaptation to perturbations in environment can be achieved by dynamically balanced interactions among consortia members. Cooperation (or mutualism) is common in consortia and is shown to be efficient for promoting biomass [17] and helpful for excluding cheaters and invaders [18]. Competition (or antagonism) is also an important and nonnegligible interaction since cells compete for space and nutrients and may associate with antibiotics warfare [19]. Work by Coyte *et al*. [20] indicate that hosts benefit from competition and stability is increased when competition dampens cooperative networks among species. Kelsic et *al*. [21] also highlights the importance of antagonism in stabilizing community structures in three-way microbial interactions.

However, most of the studies do not consider the spatial structuring of the consortia and lack theoretical explanations. Therefore, it is implicit why cooperation or competition can maintain stable coexistence in general. Spatial structures of the consortia can play a big role in population level behaviors, such as biofilm formation and cell differentiation, so realizing stable coexistence under different spatial conditions is important for consortia design.

We consider two common spatial conditions, the well-mixed scenario where no spatial information is involved for cell-cell interaction and the 2D spatial scenario where we assume cells are cultured on agar plates and perform self-organized structures. By comparing the population dynamics of cooperative and competitive interaction systems for both nonspatial and spatial scenarios, we show the conditions on achieving stable coexistence.

In Section II, we introduce a two-species interaction system of *E. coli* and demonstrate the biological design for population growth control and intra/interspecies interactions. In Section III and IV, we construct nonspatial and spatial models and show coexistence stability by simulation and linear stability analysis. In Section V, we summarize and discuss control strategies for synthetic consortia with stable coexistence.

## II. BIOLOGICAL DESIGN

We consider a two-species interaction system of *E. coli*, where *Cell*_1_ and *Cell*_2_ can interact and regulate population growth. As shown in Fig. 1(a), we denote intraspecies interaction strengths as *a_ii_*, *i* = 1,2 and interspecies interaction strengths as *a_ij_*, *i* ≠ *j*. In both cell species, there is constant production of quorum sensing signals *S*_1_ and *S*_2_. The concentrations of these small and diffusive signaling molecules can represent the population sizes of two cell species. Signaling molecules can bind with receptors to activate or repress cell growth and death processes. Mechanisms of regulating population size include toxin-antitoxin systems [22], [23], metabolite feeding [24], RNA antisense of growth gene [25] and other mechanisms. Cooperative and competitive interactions are identified by the positive and negative impacts on cell population increase via quorum sensing communication and cell growth and death actuation. For example, *Cell*_1_ produces quorum sensing signal *S*_1_. *S*_1_ activates toxin production in *Cell*_1_ that leads to death and antitoxin production in *Cell*_2_ for rescue. The intraspeceis interaction in *Cell*_1_ is competitive, i.e., *a*_11_ < 0 and the interspecies interaction from *Cell*_1_ to *Cell*_2_ is cooperative, i.e., *a*_21_ > 0.

**Fig. 1.**
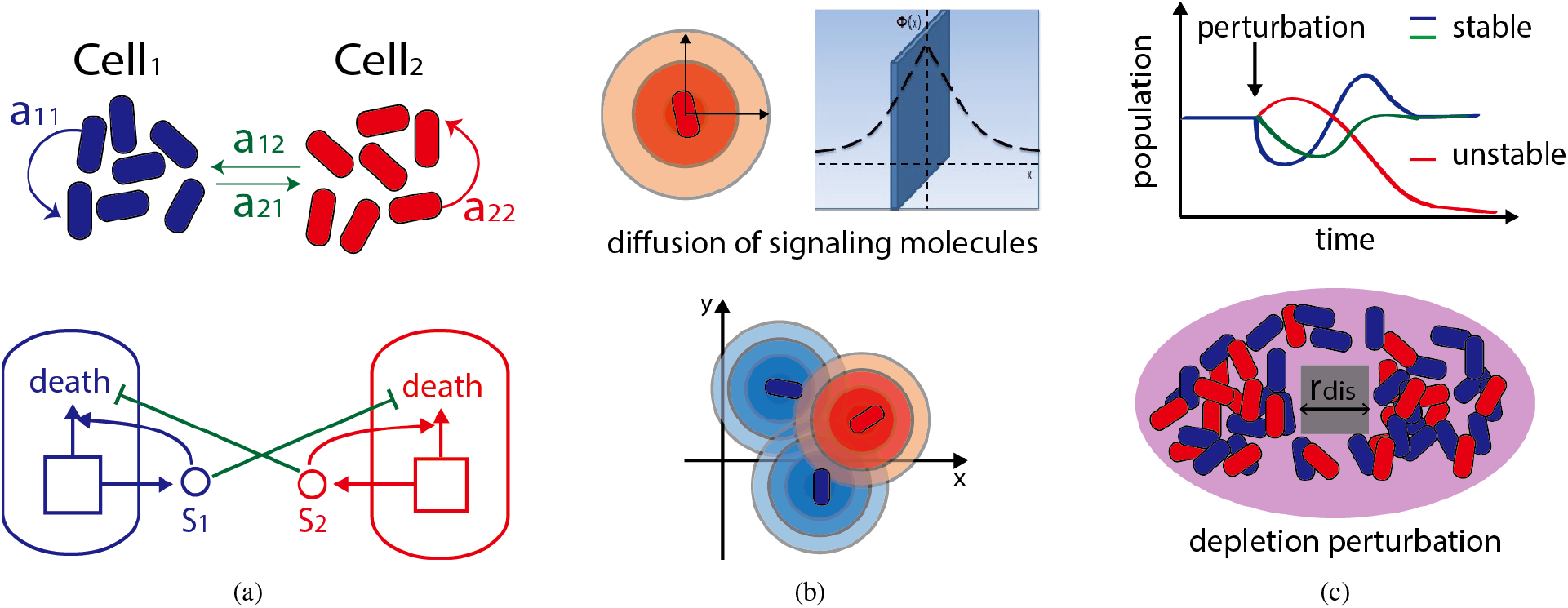
Abstract biological design and spatial effect on cell population. (a) Intra/interspecies interactions in a two-species system. The lower plot shows a design of cooperation using toxin-antitoxin mechanism. Two cell population communicate via quorum sensing signals *S*_1_,*S*_2_. In *Cell*_1_, *S*_1_ activates toxin *T* and *S*_2_ activates antitoxin production to further regulate cell death and rescue processes. Similar reactions occur in *Cell*_2_. (b) The diffusion of quorum sensing signals on an agar plate. Quorum sensing signals can diffuse around and accumulate in the environment to form a certain concentration distribution on 2D space. (c) The measurement of coexistence stability. After perturbations on cell population, e.g., antibiotics depletion within an area, if the cell population recovers, the system is stable; otherwise, it is unstable.

Cell-cell interactions depend on signaling molecules diffusing and reaching cells, and quorum sensing signals form a concentration distribution due to the heterogeneity of cell population. Therefore, it makes a big difference if the spatial condition is considered. The range of length and time scale can be affected by spatial settings of cell culture and circuit properties [26], [27]. Fig. 1(b) demonstrates that each single cell is a point source and the concentration of signaling molecules decreases when it is further from the source and leads to weaker interaction strength to other cells.

To measure the stability of a consortium, we can perturb one population in the coexisting multi-species community by antibiotics depletion or population dilution and see if the steady states recover. As shown in Fig. 1(c), if the population returns to the previous steady state, the coexistence is stable. Otherwise, the coexistence is unstable because the perturbed population goes extinct and the other species becomes dominant. Theoretically, we can assess the mathematical model and analyze the eigenvalues after linearization for local stability.

## III. NONSPATIAL MODEL AND STABILITY ANALYSIS

The basis for our population growth model is the Lotka-Volterra model. The nonspatial model represents the well-mixed scenario where every individual cell interacts with each other with identical strength and we can obtain the population interaction ODEs of two species as follows:

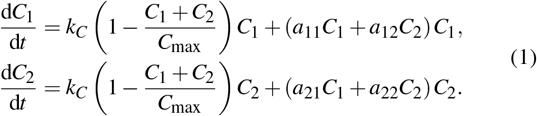

In equation (1), *k_C_* is the growth rate, *C*_max_ is the environmental carrying capacity and *a_ij_*, *i*, *j* = 1,2 are interaction strengths. The interspecies interaction is competition if *a_ij_* < 0, *i* ≠ *j*, cooperation if *a_ij_* > 0, *i* ≠ *j* and neutralism if *a_ij_* = 0, *i* ≠ *j*. To avoid growth explosion and focus on interspecies interaction properties, we only consider the scenario where intraspecies interaction is competitive, i.e., *a_ii_* < 0, *i* = 1,2. For simplicity, we assume all signaling molecules are produced and diffuse at the same rate and have same strength on activation or repression on population growth for both species, then we have *a*_11_ = *a*_22_,*a*_12_ = *a*_21_. In general, the interaction strengths may be asymmetrical for two species, and the steady states of population size are not ideally at 1:1 ratio. The following analysis methods still apply.

There exist three nonzero steady states for equation (1). One steady state is a nontrivial solution where both species exist, denoted as 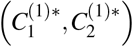 and the other two steady states indicate one species dominates and the other species dies, denoted as 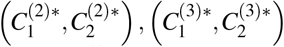. We can solve for the steady states as follows:

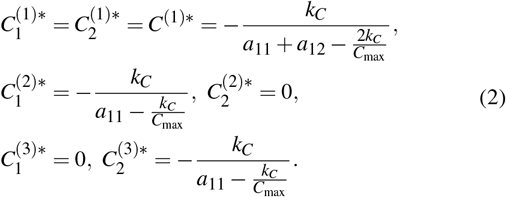

The equilibrium 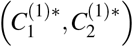 corresponds to coexistence of two cell populations. We set parameters |*a*_12_| = 0.8|*a*_11_| for both cooperative and competitive interactions and rescale the parameter values to achieve the same equilibrium at *C*^(1)^* = 100. We set initial conditions to be random nonzero values. In simulations, we perturb the system by diluting out 20%, 40%, 60% of *Cell*_1_ population after reaching steady state at 1:1 ratio at *t* = 50 hr, and measure the recovery. Using bioSCRAPE toolbox for deterministic and stochastic simulations [28], we show that cooperation and competition maintain stable coexistence in Fig. 2, where the relative population ratios between *Cell*_1_ and *Cell*_2_ all recover to the 1:1 ratio.

**Fig. 2.**
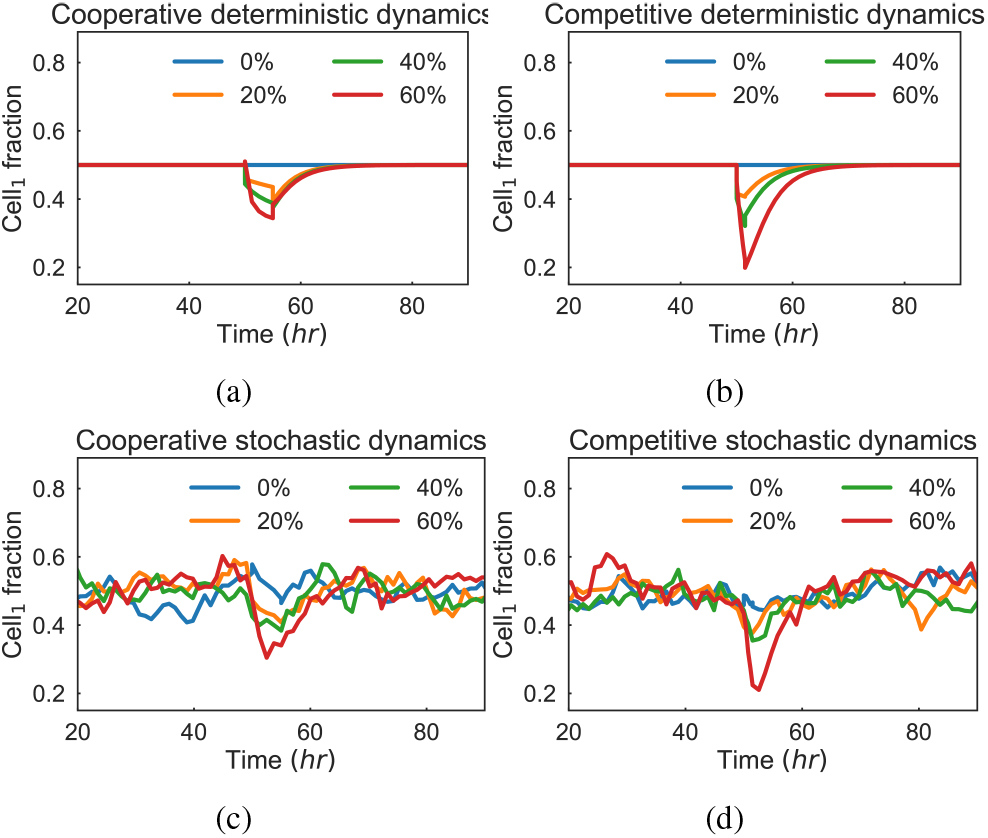
Simulations of cell population dynamics under perturbation for nonspatial models. Panels (a) and (b) are the deterministic simulations for cooperative and competitive interactions under different levels of cell depletion on *Cell*_1_. Panels (c) and (d) are the stochastic simulations for cooperative and competitive interactions under different levels of cell depletion on *Cell*_1_.

We next investigate stability conditions for both interactions from the nonspatial model. We first analyze the local stability at equilibrium 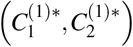 by linearizing equation (1) as

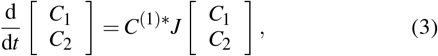

where the Jacobian matrix is defined as

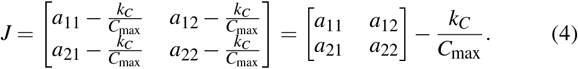

According to local stability criteria and *C*^(1)^* > 0, this requires

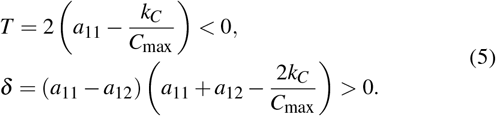

Equation (5) is equivalent to

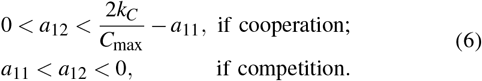

Fig. 3 shows the region of stable and unstable coexistence by identifying dominant eigenvalues with negative and non-negative real parts when altering interaction strengths. Constraints on *a*_12_ in equation (6) confirm that as long as the interspecies interaction strength is weaker than the intraspecies interaction, the consortia is stable for both cooperation and competition. It also matches with the simulation results in Fig. 2 since we set the parameters to have |*a*_12_| < |*a*_11_| and *a*_11_ < 0.

**Fig. 3.**
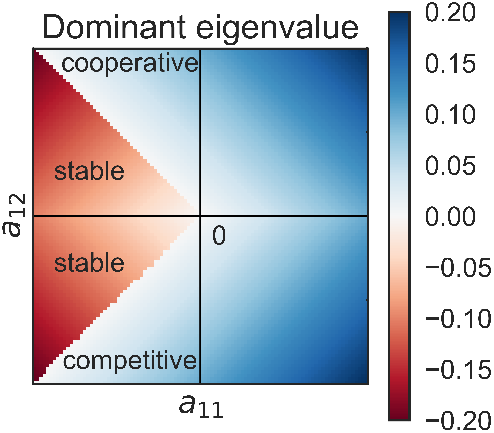
Stability conditions at the coexistence equilibrium for the nonspatial interaction model. Dominant eigenvalues are calculated under different sets of intra/interspecies interaction strengths *a*_11_ and *a*_12_. If the real parts of eigenvalues are negative, shown in red color, the coexistence is stable. Otherwise, the coexistence is unstable, as shown in blue. Both cooperative and competitive interactions have large stable regimes.

We show the coexistence is globally stable by introducing the following theorem from [29].

### Theorem 1.

If the nontrivial equilibrium of equation (1) is feasible and there exist a constant positive diagonal matrix P such that PJ + J^T^P is negative definite, then equation (1) is globally stable in the feasible region.

Here our assumption of *a*_11_ = *a*_22_,*a*_12_ = *a*_21_ ensures that *J* is already negative definite when equation (6) is satisfied. For more general cases when interaction strengths are not symmetrical for both species, it is easy to find a constant positive diagonal matrix *P* in such form:

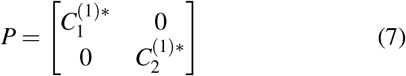

that satisfies *Theorem* 1 and derive global stability from local stability.

## IV. SPATIAL MODEL WITH DIFFUSION AND NONLOCAL REACTIONS

Under the assumption that cells can grow and move to access more space and resources, it is natural to model the spatial condition as a reaction-diffusion system using PDEs. Diffusion is introduced in [30] to describe cell motility in the spatial environment, in forms of Δ (*D* (*f* (*u*, *v*)) *u*) where *u*, *v* are cell population densities and 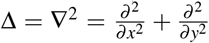 represents the Laplace operator. The specific function *f* (*u*, *v*) depends on cell-cell interactions. In our design, we assume that cells diffuse faster if the the growth is activated since growth factors can increase cell division and movement [31], and the function *f* (*u*, *v*) can be characterized by intra/interspecies interactions. All intra/interspecies interactions are realized via quorum sensing and the diffusible signaling molecules can only reach cells in the neighborhood within some range. Therefore, the interactions are nonlocal behaviors that depend on the spatial distribution of cells and of signaling molecules in the neighborhood. We assume cells are point sources of diffusible signaling molecules on 2D space. Adding source production and self decay in signaling molecules diffusion equations, we derive the signaling molecules concentration *ϕ* at the radius *r* of a single source at steady state with appropriate boundary conditions as

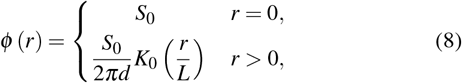

where *S*_0_ is the production rate of signaling molecules, *d* is the diffusion rate of signaling molecules, *L* is the diffusion range calculated as 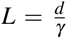, *γ* is the degradation rate of signaling molecules and *K*_0_ is the modified Bessel function of the second kind of order zero, which can be approximated as the inverse of a log function when *r* is small. Therefore, the interactions are no longer linear functions of cell population densities as in equation (1), but instead are weighted by a decreasing distance kernel *ϕ* (*r*). To be consistent with the parameters in the nonspatial model, we let

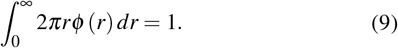

The interactions on cells at position ***x*** ∈ ℝ^2^ are nonlocal in the following form:

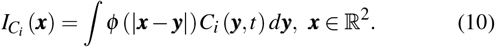

Such nonlocal interactions also have an impact on cell diffusion dynamics in the following manner, since cells divide and move faster where there are more cooperative than competitive interaction signals:

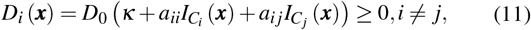

where *κ* is a basal diffusion scaled by *D*_0_. Note that the competitive and cooperative interactions affect the spatial system via cell population growth and cell movement consistently. By substituting *C_i_*, *i* = 1,2 in interaction terms with equation (10) and adding cell diffusion according to equation (11), we extend the nonspatial ODE model in equation (1) into a PDE model as

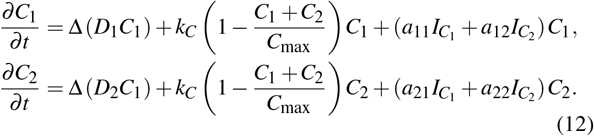

We solve for homogeneous steady states of equation (12) and obtain three nonzero solutions coincident with equation (2). The nontrivial homogeneous solution is *C*_1_ (***x***, *t*) = *C*_2_ (***x***, *t*) ≡ *C*^(1)^*.

We use same parameters and run simulations in gro [32]. We set the initial condition as a homogeneous distribution and observe self-organized spatial structures of two species. Both cooperation and competition maintain coexistence after 120 hr. As shown in Fig. 4(a) and 4(b), cooperation leads to a more intermixed spatial pattern while competition tends to self-organize into small patches of segregated colonies. When we deplete some of *Cell*_1_ population within range rdis at *t* = 50 hr, cooperation helps recovery of *Cell*_1_ population but competition lets *Cell*_2_ dominates the antibiotics dispersal area and extincts *Cell*_1_ completely. Fig. 4(c) and 4(d) show the population fraction dynamics when rdis is altered and only cooperation maintains stable coexistence after perturbation.

**Fig. 4.**
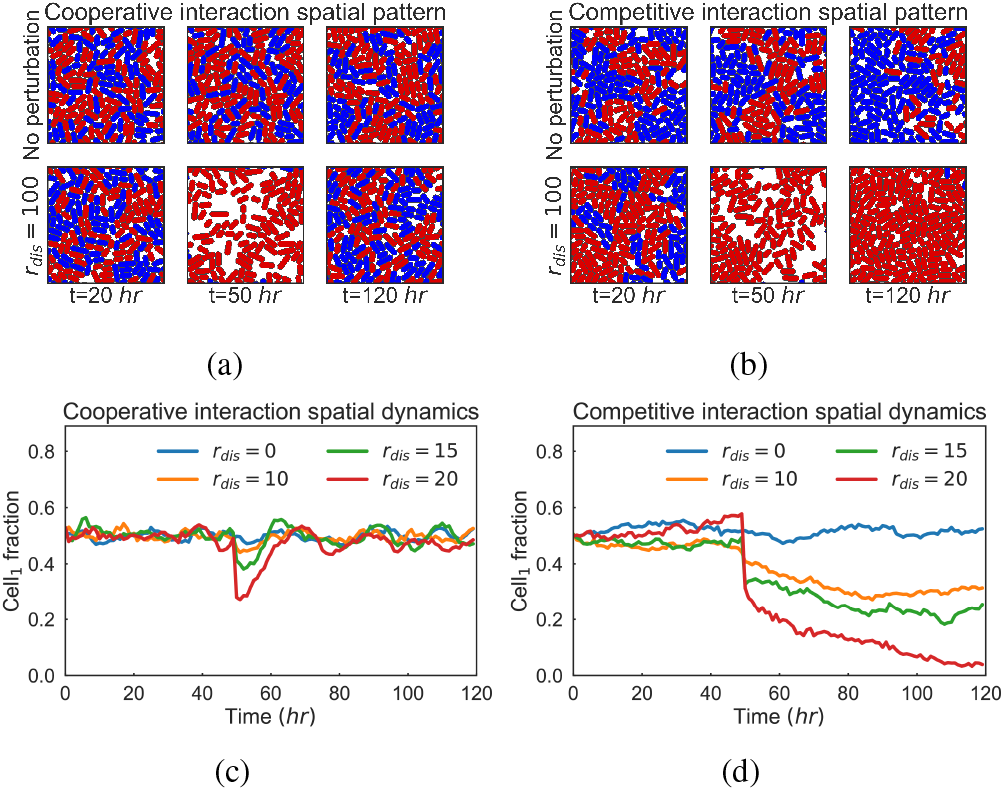
Simulations of cell population patterning and dynamics under perturbation for spatial models. Panels (a) and (b) are spatial patterns for cooperative and competitive interactions w/o perturbation on *Cell*_1_ population within an area of size *r_dis_* = 100. *Cell*_1_ is shown in blue color and *Cell*_2_ in red color in the snapshots. In (a), the upper patterns show that two cell species are forming intermixed patterns without perturbation. The lower patterns show that cooperation helps recovering coexistence and intermixed patterns after the perturbation. In (b), the upper patterns show that two cell species are forming segregated patches without perturbation. The lower patterns show that the coexistence breaks as *Cell*_2_ dominates the perturbed area and forms a big segregation. Panels (c) and (d) are corresponding population dynamics for cooperation and competition with different perturbation area sizes.

Now we give theoretical explanations of the significant difference of coexistence stability under spatial conditions between cooperation and competition. For a coexisting homogeneous steady state *C*_1_ (*x*, *t*) = *C*_2_ (*x*, *t*) ≡ *C*^(1)^* > 0, we linearize equation (12) around equilibrium, apply Fourier transform and obtain the following characteristic equation given *C*^(1)^* > 0:

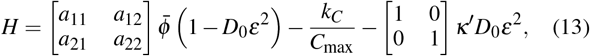

where 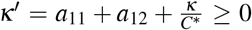. The Fourier transform of *ϕ* is 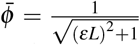 according to equation (8) and (9). The local stability of the homogeneous coexistence under spatial perturbation requires the following conditions,

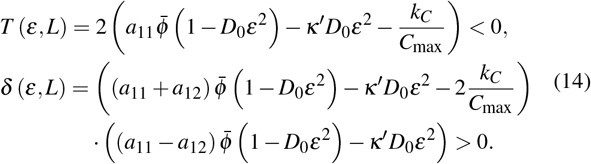

Equation (14) is equivalent to equation (5) for the nonspatial model when there is no cell and signaling molecule diffusion. In other words, nonspatial conditions can be described by equation (14) when *L* = 0 and *D*_0_ = 0. However, given parameters satisfying nonspatial stability conditions, the local stability of homogeneous solution is different for cooperative and competitive interactions.

In Fig. 5, we alter cell population diffusion rate *D*_0_ and quorum sensing signaling molecules diffusion range *L*, and calculate *T* and δ to identify stable regions. For cooperative interactions, the homogeneous coexistence is stable as equation (14) is satisfied and populations form intermixed spatial structures. When *Cell*_1_ in certain area is perturbed, cells outside this area could diffuse in and the other existing species *Cell*_2_ would activate *Cell*_1_’s growth to recover coexistence. Similar beneficial phenomena have been observed where cooperative partners spatially intermix by appearing successively on top of each other in engineered metabolites exchanging consortia [33].

**Fig. 5.**
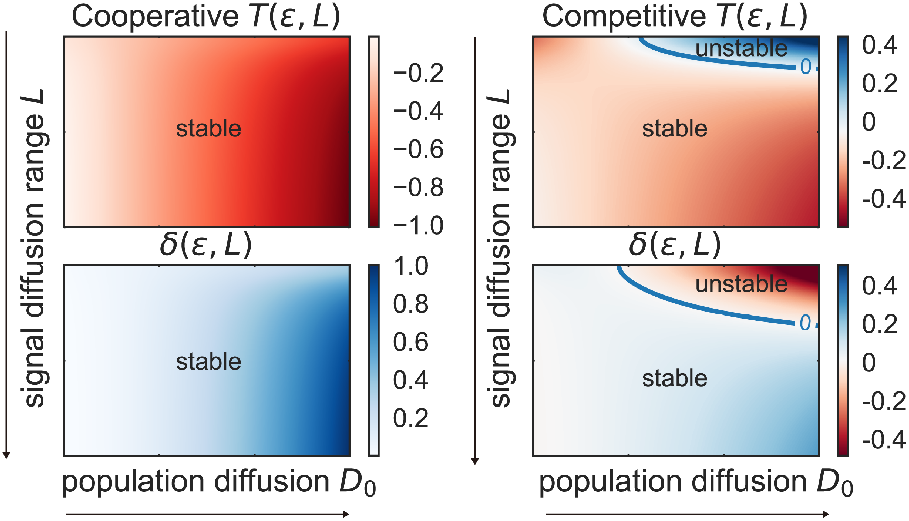
Stability conditions of homogeneous coexistence for the spatial interaction model. We calculate *T* in the upper plots and δ in the lower plots for two interactions when altering two parameters *D*_0_ and *L*. Given the stability conditions in equation (14), the homogeneous coexistence is stable in parameter regimes where *T* < 0 as shown in red color and δ > 0 as shown in blue color. The left plots show that cooperation maintains stable homogeneous coexistence. The right plots show that it can be unstable for competitive interaction at high population diffusion rate and small quorum sensing signal diffusion range.

For competitive interactions, the stability condition may not be satisfied for some parameter regimes. Large population diffusion rates indicate the strong repression on cell motility from competitive interactions and small signal diffusion ranges stress the impact of spatial heterogeneity of signaling molecules on population dynamics. Thus, spatial perturbations can break the coexistence stability when *D*_0_ increases and *L* is small. Since two cell species repress the population growth of each other, once *Cell*_1_ senses *Cell*_2_ population, it dies more until the decrease in population compensates for self repression. It is harder for cells to mix but instead they form small segregated colonies. When *Cell*_1_ are depleted, only *Cell*_2_ grows and prevents *Cell*_1_ from diffusing into the area, and eventually outcompetes *Cell*_1_. Thus, one dominant cell population cannot recover coexistence after the perturbation. Spatial perturbations on cell population are common noises in environment, therefore cooperative interaction is more robust than competitive interaction in consortia.

## V. DISCUSSION

In this paper, we show that cooperation leads to intermixed patterns and supports more stable coexistence so that it is more robust to population perturbations under spatial condition. Competition is easily perturbed and becomes unstable for coexistence. When cells are well-mixed, both interactions are stable, but when cells grow and self-organize into certain spatial structures, the nonlocal interaction behavior and population diffusion are important factors that cause such different stability performance between cooperation and competition. The results can provide useful guidance when we engineer synthetic consortia, especially when strong spatial heteroeneity in environment is considered. Cooperative interaction is beneficial because it maintains stable coexistence, while competition can better perform localized functions by forming spatially segregated colonies. For example, the human microbiome consists of hundreds of microbial species and they are grouped to perform complicated functions [34]. It is important to keep stable and robust coexistence within groups and avoid cross-talk and interference from unrelated groups at the same time. By implementing certain interspecies interaction type and regulating the interaction strength or signal diffusion range, we can improve the the performance of synthetic consortia dealing with multiple tasks. Another application could be spatial-temporal control on cell differentiation. The rate of evolution depends on spatial organization and interactions among the population in nontrivial ways. It has been shown that cooperation leads to fast creation of complex phenotypes as an emergent property [35].

Our work is based on a general interaction model of two-species systems, but the analysis can be applied to more circuits and problems in synthetic biology. Different biological mechanisms can be implemented to perform population level functions with theoretical predictions. For future work, we would like to construct specific synthetic consortia using potential cooperative and competitive interaction control on population regulation and explore the spatial effect on coexistence stability and robustness to perturbations in consortia containing more species and complex interaction networks.

## VI. ACKNOWLEDGMENTS

The authors would like to thank Fangzhou Xiao for his insightful discussion. The author X. R is partially supported by the Air Force Office of Scientific Research, grant number FA9550-14-1-0060. The project depicted is also sponsored by the Defense Advanced Research Projects Agency (Agreement HR0011-17-2-0008). The content of the information does not necessarily reflect the position or the policy of the Government, and no official endorsement should be inferred.

## REFERENCES

[1] J. E. Koenig, A. Spor, N. Scalfone, A. D. Fricker, J. Stombaugh, R. Knight, L. T. Angenent, and R. E. Ley, “Succession of microbial consortia in the developing infant gut microbiome,” Proceedings of the National Academy of Sciences, vol. 108, no. 1, pp. 4578–4585, 2011.

[2] L. Cocolin and D. Ercolini, “Zooming into food-associated microbial consortia: a ‘cultural’evolution,” Current Opinion in Food Science, vol. 2, pp. 43–50, 2015.

[3] B. K. Singh, P. Millard, A. S. Whiteley, and J. C. Murrell, “Unravelling rhizosphere-microbial interactions: opportunities and limitations,” Trends in Microbiology, vol. 12, no. 8, pp. 386–393, 2004.

[4] E. A. Bayer, R. Lamed, and M. E. Himmel, “The potential of cellulases and cellulosomes for cellulosic waste management,” Current Opinion in Biotechnology, vol. 18, no. 3, pp. 237–245, 2007.

[5] K. Brenner, L. You, and F. H. Arnold, “Engineering microbial consortia: a new frontier in synthetic biology,” Trends in Biotechnology, vol. 26, no. 9, pp. 483–489, 2008.

[6] W. Shou, S. Ram, and J. M. Vilar, “Synthetic cooperation in engineered yeast populations,” Proceedings of the National Academy of Sciences, vol. 104, no. 6, pp. 1877–1882, 2007.

[7] S. R. Lindemann, H. C. Bernstein, H.-S. Song, J. K. Fredrickson, M. W. Fields, W. Shou, D. R. Johnson, and A. S. Beliaev, “Engineering microbial consortia for controllable outputs,” The ISME Journal, vol. 10, no. 9, p. 2077, 2016.

[8] F. K. Balagaddé, H. Song, J. Ozaki, C. H. Collins, M. Barnet, F. H. Arnold, S. R. Quake, and L. You, “A synthetic escherichia coli predator-prey ecosystem,” Molecular Systems Biology, vol. 4, no. 1, 2008.

[9] J. R. Nahum, B. N. Harding, and B. Kerr, “Evolution of restraint in a structured rock-paper-scissors community,” Proceedings of the National Academy of Sciences, vol. 108, no. Supplement 2, pp. 10831–10838, 2011.

[10] Y. Chen, J. K. Kim, A. J. Hirning, K. Josic, and M. R. Bennett, “Emergent genetic oscillations in a synthetic microbial consortium,” Science, vol. 349, no. 6251, pp. 986–989, 2015.

[11] M. Sadeghpour, A. Veliz-Cuba, G. Orosz, K. Josić, and M. R. Bennett, “Bistability and oscillations in co-repressive synthetic microbial consortia,” Quantitative Biology, vol. 5, no. 1, pp. 55–66, 2017.

[12] G. Fiore, A. Matyjaszkiewicz, F. Annunziata, C. Grierson, N. J. Savery, L. Marucci, and M. di Bernardo, “Design of a multicellular feedback control strategy in a synthetic bacterial consortium,” in 2016 IEEE 55th Conference on Decision and Control (CDC), 2016, pp. 3338–3343.

[13] W. Kong, D. R. Meldgin, J. J. Collins, and T. Lu, “Designing microbial consortia with defined social interactions,” Nature Chemical Biology, p. 1, 2018.

[14] M. B. Miller and B. L. Bassler, “Quorum sensing in bacteria,” Annual Reviews in Microbiology, vol. 55, no. 1, pp. 165–199, 2001.

[15] D.-K. Ro, E. M. Paradise, M. Ouellet, K. J. Fisher, K. L. Newman, J. M. Ndungu, K. A. Ho, R. A. Eachus, T. S. Ham, J. Kirby, M. C. Y. Chang, S. T. Withers, Y. Shiba, R. Sarpong, and J. D. Keasling, “Production of the antimalarial drug precursor artemisinic acid in engineered yeast,” Nature, vol. 440, no. 7086, pp. 940–943, 2006.

[16] T. Arai, S. Matsuoka, H.-Y. Cho, H. Yukawa, M. Inui, S.-L. Wong, and R. H. Doi, “Synthesis of clostridium cellulovorans minicellulosomes by intercellular complementation,” Proceedings of the National Academy of Sciences, vol. 104, no. 5, pp. 1456–1460, 2007.

[17] M. Burmølle, J. S. Webb, D. Rao, L. H. Hansen, S. J. Sørensen, and S. Kjelleberg, “Enhanced biofilm formation and increased resistance to antimicrobial agents and bacterial invasion are caused by synergistic interactions in multispecies biofilms,” Applied and Environmental Microbiology, vol. 72, no. 6, pp. 3916–3923, 2006.

[18] S. Pande, F. Kaftan, S. Lang, A. Svatoš, S. Germerodt, and C. Kost, “Privatization of cooperative benefits stabilizes mutualistic cross-feeding interactions in spatially structured environments,” The ISME Journal, vol. 10, no. 6, p. 1413, 2016.

[19] J. C. Clemente, E. C. Pehrsson, M. J. Blaser, K. Sandhu, Z. Gao, B. Wang, M. Magris, G. Hidalgo, M. Contreras, Ó. Noya-Alarcón, et al., “The microbiome of uncontacted amerindians,” Science advances, vol. 1, no. 3, p. e1500183, 2015.

[20] K. Z. Coyte, J. Schluter, and K. R. Foster, “The ecology of the microbiome: networks, competition, and stability,” Science, vol. 350, no. 6261, pp. 663–666, 2015.

[21] E. D. Kelsic, J. Zhao, K. Vetsigian, and R. Kishony, “Counteraction of antibiotic production and degradation stabilizes microbial communities,” Nature, vol. 521, no. 7553, p. 516, 2015.

[22] L. You, R. S. Cox, R. Weiss, and F. H. Arnold, “Programmed population control by cell-cell communication and regulated killing,” Nature, vol. 428, no. 6985, pp. 868–871, 2004.

[23] R. D. McCardell, S. Huang, L. N. Green, and R. M. Murray, “Control of bacterial population density with population feedback and molecular sequestration,” bioRxiv, p. 225045, 2017.

[24] J. J. Bull and W. R. Harcombe, “Population dynamics constrain the cooperative evolution of cross-feeding,” PLoS One, vol. 4, no. 1, p. e4115, 2009.

[25] X. Ren, A.-A. Baetica, A. Swaminathan, and R. M. Murray, “Population regulation in microbial consortia using dual feedback control,” in Decision and Control (CDC), 2017 IEEE 56th Annual Conference on. IEEE, 2017, pp. 5341–5347.

[26] G. E. Dilanji, J. B. Langebrake, P. De Leenheer, and S. J. Hagen, “Quorum activation at a distance: spatiotemporal patterns of gene regulation from diffusion of an autoinducer signal,” Journal of the American Chemical Society, vol. 134, no. 12, pp. 5618–5626, 2012.

[27] J. Doong, J. Parkin, and R. M. Murray, “Length and time scales of cell-cell signaling circuits in agar,” bioRxiv, p. 220244, 2017.

[28] A. Swaminathan, V. Hsiao, and R. M. Murray, “Quantitative modeling of integrase dynamics using a novel python toolbox for parameter inference in synthetic biology,” 2017.

[29] B. S. Goh, “Global stability in many-species systems,” The American Naturalist, vol. 111, no. 977, pp. 135–143, 1977.

[30] K. J. Painter and J. A. Sherratt, “Modelling the movement of interacting cell populations,” Journal of theoretical biology, vol. 225, no. 3, pp. 327–339, 2003.

[31] B. Anand-Apte and B. Zetter, “Signaling mechanisms in growth factor-stimulated cell motility,” Stem Cells, vol. 15, no. 4, pp. 259–267, 1997.

[32] S. S. Jang, K. T. Oishi, R. G. Egbert, and E. Klavins, “Specification and simulation of synthetic multicelled behaviors,” ACS Synthetic Biology, vol. 1, no. 8, pp. 365–374, 2012.

[33] B. Momeni, K. A. Brileya, M. W. Fields, and W. Shou, “Strong inter-population cooperation leads to partner intermixing in microbial communities,” Elife, vol. 2, p. e00230, 2013.

[34] C. Tropini, K. A. Earle, K. C. Huang, and J. L. Sonnenburg, “The gut microbiome: connecting spatial organization to function,” Cell Host & Microbe, vol. 21, no. 4, pp. 433–442, 2017.

[35] N. L. Komarova, “Spatial interactions and cooperation can change the speed of evolution of complex phenotypes,” Proceedings of the National Academy of Sciences, vol. 111, no. Supplement 3, pp. 10789–10795, 2014.

